# Estimating fMRI Timescale Maps

**DOI:** 10.1101/2025.04.23.650300

**Authors:** Gabriel Riegner, Samuel Davenport, Bradley Voytek, Armin Schwartzman

## Abstract

Brain activity unfolds over hierarchical timescales that reflect how brain regions integrate and process information, linking functional and structural organization. While timescale studies are prevalent, existing estimation methods rely on the restrictive assumption of exponentially decaying temporal autocorrelation and only provide point estimates without standard errors, limiting statistical inference. In this paper, we formalize and evaluate two methods for mapping timescales in resting-state fMRI: a time-domain fit of an autoregressive (AR1) model and an autocorrelation-domain fit of an exponential decay model. Rather than assuming exponential autocorrelation decay, we define timescales by projecting the fMRI time series onto these approximating models, requiring only stationarity and mixing conditions while incorporating robust standard errors to account for model misspecification. We introduce theoretical properties of timescale estimators and show parameter recovery in realistic simulations, as well as applications to fMRI from the Human Connectome Project. Comparatively, the time-domain method produces more accurate estimates under model misspecification, remains computationally efficient for high-dimensional fMRI data, and yields maps aligned with known functional brain organization. In this work, we show valid statistical inference on fMRI timescale maps, and provide Python implementations of all methods.

## 1 Introduction

### 1.1 fMRI Timescale Maps

Neural processes span multiple timescales, from millisecond synaptic events to slower activity coordinating distributed brain networks (Buzsáki, 2004). Timescales index the integration of neural activity over time, defined by exponential decay in autocorrelation that emerges from stochastic models of neural activity (Murray et al., 2014, Supplementary Mathematical Note). When mapped across the cortex, these local temporal timescales reveal a large-scale spatial organization. Multimodal evidence links these timescale differences to intrinsic brain hierarchies, reflecting how distinct regions integrate and process information over time. Timescale maps of the brain align with functional hierarchy – sensory areas that process rapidly changing stimuli show shorter timescales than association areas involved in cognitive processes that unfold over longer durations (Raut et al., 2020; Gao et al., 2020; Hasson et al., 2008; Murray et al., 2014; Stephens et al., 2013). This hierarchy is also associated with anatomical organization including myelination levels and gene expression patterns, as shown in studies using human electrophysiology, MEG, and gene expression profiling (Gao et al., 2020; Shafiei et al., 2023).

Computational modeling by Li and Wang (2022) suggests that hierarchical timescales emerge from: (i) brain-wide gradients in synaptic excitation strength, (ii) electrophysiological differences between excitatory and inhibitory neurons, and (iii) balance between distant excitatory and local inhibitory inputs. In addition to these intrinsic mechanisms, there is growing evidence that neuronal timescales are dynamic and modulated by experimental manipulations or behavioral demands. For example, pharmacological agents like propofol and serotonergic drugs alter intrinsic timescales, affecting the temporal integration of information in the brain (Huang et al., 2018; Shinn et al., 2023). Timescale changes have also been observed during development, sleep deprivation, wakefulness, neuropsychiatric disorders (autism and schizophrenia), and naturalistic behaviors (Martin-Burgos et al., 2024; Meisel et al., 2017; Watanabe et al., 2019; Wengler et al., 2020; Manea et al., 2024). These findings demonstrate that timescales are broadly relevant to both structural and functional properties of the brain.

Seminal research on timescales has primarily used invasive electrophysiology in non-human animals (Murray et al., 2014; Cirillo et al., 2018; Nougaret et al., 2021; Manea et al., 2022; Spitmaan et al., 2020; Trepka et al., 2024). While these methods provide high temporal resolution for studying neural activity up to the single-neuron level, they are limited by sparse spatial sampling. Investigating the large-scale spatial organization of timescale maps requires non-invasive methods like resting-state functional MRI (rfMRI), which measures spontaneous fluctuations in the blood oxygen level-dependent (BOLD) signal. Unlike techniques with sparser spatial coverage such as EEG or MEG, rfMRI provides dense, whole-brain coverage, though its power is concentrated in the much slower (<0.1 Hz) frequency range (Raut et al., 2020; He, 2011). This is because the BOLD signal reflects hemodynamic changes, which act as a low-pass filter on neural activity, and because the signal-to-noise ratio is highest in this infraslow frequency band. Although an indirect measure, the BOLD signal reflects functionally significant fluctuations associated with underlying electrophysiological activity (Logothetis, 2008), making it a valuable tool for investigating high spatial-resolution cortical timescale maps. Studies have shown that rfMRI-derived timescale maps align spatially with those from other imaging modalities across human and animal models (Raut et al., 2020; Shafiei et al., 2020; Lurie et al., 2024). The present study will focus on rfMRI from the Human Connectome Project dataset (Van Essen et al., 2013).

### 1.2 Current methods

A key consideration is the representation of the time series – whether in its original time domain, or transformed into another domain for analysis. Timescales are generally estimated using three main methods corresponding to neural data represented in the (i) time domain, (ii) autocorrelation domain, or (iii) frequency domain. The most common is the autocorrelation domain, where timescales are defined by fitting an exponential decay model to the sample autocorrelation function (ACF) (Rossi-Pool et al., 2021; Cirillo et al., 2018; Ito et al., 2020; Runyan et al., 2017; Zeraati et al., 2022; Nougaret et al., 2021; Wasmuht et al., 2018; Müller et al., 2020; Maisson et al., 2021; Li and Wang, 2022; Shafiei et al., 2020). Similar approaches compute timescales directly from the sample ACF as the sum of positive autocorrelations (Wengler et al., 2020; Manea et al., 2022; Watanabe et al., 2019), or by identifying where the sample ACF crosses a specified threshold (Wengler et al., 2020; Zilio et al., 2021). Alternatively, the time-domain method uses a first-order autoregressive (AR1) model to estimate timescales directly from time series (Kaneoke et al., 2012; Meisel et al., 2017; Huang et al., 2018; Lurie et al., 2024; Shinn et al., 2023; Shafiei et al., 2020; Spitmaan et al., 2020; Trepka et al., 2024), and has shown better test-retest reliability than autocorrelation-domain methods for rfMRI (Huang et al., 2018). Finally, frequency-domain methods operate on the premise that timescales are properties of the brain’s aperiodic (non-rhythmic) activity. These methods therefore use the power-spectral density to estimate and remove oscillatory components to mitigate their influence on timescale estimates (Donoghue et al., 2020; Gao et al., 2020; Manea et al., 2024; Zeraati et al., 2022; Fallon et al., 2020). This paper focuses on time- and autocorrelation-domain methods, avoiding external criteria that attempt to separate periodic from aperiodic contributions to the BOLD signal. Because amplitude-modulated rhythmic activity can mimic aperiodic properties (He et al., 2010; He, 2011), distinguishing the true generative process is often circular. Instead, we introduce a timescale estimator that is robust regardless of whether the underlying process is strictly aperiodic, damped-oscillatory, or a mixture of both, and we validate this robustness using simulations across these regimes.

### 1.3 Problem Statement and Proposed Solution

A key challenge in applied timescale research is the lack of standardized model definitions, where diverse approaches have led to inconsistent findings across studies (Zeraati et al., 2022; Fallon et al., 2020; Shafiei et al., 2020). Many parameterization methods rely on restrictive assumptions, such as exponential autocorrelation decay, which may bias timescale estimates and (more often) their standard errors (Woolrich et al., 2001; Zeraati et al., 2022; Raut et al., 2019). Additionally, the distributional properties of these methods are often ignored, resulting in studies reporting only point estimates without quantifying uncertainty, which limits statistical inference and hypothesis testing.

To address these issues, this paper formalizes and evaluates two commonly applied timescale methods in rfMRI: the time-domain fit of an autoregressive (AR1) model and the autocorrelation-domain fit of an exponential decay model. The goal is to estimate accurate timescale maps that enable robust statistical testing and inference across brain regions. This work offers the following contributions: (i) The assumptions are generalized to include processes that are both stationary *and* mixing, not only those with exponential autocorrelation decay. (ii) Robust standard errors account for the inevitable mismatch between the data-generating process and fitted model, enabling valid inference despite model misspecification. (iii) Theoretical properties demonstrate that both time- and autocorrelation-domain estimators converge to different values due to their distinct definitions, and are consistent (i.e., converge in probability to the method-specific estimand) and asymptotically normal. (iv) Simulations confirm that both methods yield unbiased estimates across autoregressive and realistic settings, with standard errors that are robust to non-exponential autocorrelation decay. (v) Empirical analysis of rfMRI from the Human Connectome Project shows that both approaches yield similar t-ratio maps (timescales relative to their standard errors), revealing a hierarchical organization of timescales across the cortex. While this hierarchy aligns with prior point estimate maps, our approach improves interpretability by accounting for uncertainty, ensuring that observed patterns are not artifacts of sampling noise or model misspecification. (vi) Comparative Monte Carlo simulations and empirical data analyses show that the time-domain method performs as well as, and often better than, the autocorrelation-domain method, with improved computational efficiency for high-dimensional fMRI data.

The proposed methods address important limitations in fMRI timescale research by providing rigorous statistical methods that move beyond point estimates to incorporate uncertainty quantification. This work establishes a methodological foundation for reliable inference, organized as follows. First, Section 2 formalizes the statistical models and their properties. Numbered equations indicate important expressions that are referenced throughout, while the others provide intermediate steps that can be skipped without loss of continuity. These models are validated against varying degrees of realistic misspecification in Section 3, followed by a demonstration of their empirical utility in Section 4 using the Human Connectome Project dataset. Finally, Section 5 addresses practical considerations for the interpretation of these estimates in the context of fMRI — detailing the influence of hemodynamics, measurement noise, and periodicity — to support future research investigating the functional and structural organization of timescales in the brain.

## 2 Methods

There are varying definitions and conventions in the study of time series data and stochastic processes more generally. To simplify our presentation, this work closely follows the Hansen (2022) *Econometrics* textbook as a primary reference. Throughout, we will frequently refer the reader to specific results from this textbook.

### 2.1 Assumptions

Let {*X*_*t*_} _*t* ∈ {…,−2,−1,0,1,2,…_} be a discrete-time stochastic process that is Weakly stationary and Strong Mixing (abbreviated as “stationary” and “mixing” throughout), and let {*x* = *x*_1_, *x*_2_, …, *x*_*T*_} where *x*_*t*_ denotes an observed sample of *X*_*t*_. For simplicity, assume 𝔼[*X*_*t*_] = 0.

#### 2.1.1 Weakly stationary

Weak stationarity implies a constant mean and variance (independent of time index *t*), and an autocovariance function that only depends on time lag *k, γ*_*k*_ = cov[*X*_*t*_, *X*_*t*−*k*_] = 𝔼[*X*_*t*_*X*_*t*−*k*_]. For analysis, we use a normalized measure of the autocovariances, the *autocorrelation function (ACF)*:

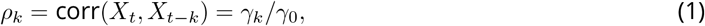

where *γ*_*k*_ is the autocovariance at lag *k* and *γ*_0_ is the variance. We emphasize that *ρ*_*k*_ represents *temporal* autocorrelation, not spatial autocorrelation often referenced in the fMRI literature. As a correlation, |*ρ*_*k*_| ≤ 1, but stationarity alone does not guarantee decay of *ρ*_*k*_ with increasing time lag; constant or periodic processes can maintain nonzero correlations indefinitely.

#### 2.1.2 Strong Mixing

Strong mixing (*α*-mixing) imposes stronger dependence constraints than stationarity while still allowing for a wide set of stochastic processes. By definition, a process is strong mixing if *α*(*𝓁*) → 0 as 𝓁 → ∞, where *α*(𝓁) measures the *maximum dependence* between events separated by 𝓁 time points. This controls the full joint dependence structure and implies distant events are uncorrelated. Strong mixing also implies ergodicity (Hansen, 2022, Chapter 14.12), which ensures consistent estimation by the ergodic theorem (Hansen, 2022, Theorem 14.9). Additionally, if a mixing process has *r >* 2 finite moments 𝔼E |*X*_*t*_| ^*r*^ < ∞and its mixing coefficients satisfy 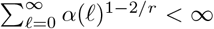, then its autocorrelations decay sufficiently fast for application of the central limit theorem for dependent data (Hansen, 2022, Theorem 14.15). These conditions enable asymptotic theory for estimation and inference on timescale maps (see Estimator Properties).

As introduced by Murray et al. (2014), the timescale *τ* represents the lag where exponentially decaying autocorrelations reach 1/*e ≈* 0.37, analogous to the time constants of many physical systems, and often referred to as *e*-folding time in other fields. While it provides an intuitive description of the memory or persistence of that process, assuming an exponential function imposes stricter constraints than strong mixing, which alone does not prescribe any specific type of decay (exponential, linear, damped periodic, etc.). Despite this, the timescale serves as a useful summary statistic for the duration of temporal dependence, enabling direct comparisons across brain regions even when the true decay structure is more complex. This highlights an important distinction between the data-generating process and the simplified parametric model used to describe the timescale at which such a process becomes decorrelated. In the present paper, we adopt broad assumptions, requiring only that the process is stationary and mixing, to account for cases where the ACF decay may not be strictly exponential.

#### 2.1.3 Estimation and Inference under Misspecification

We approximate the dominant exponential decay in autocorrelations by a single timescale parameter *τ*, and formally evaluate two timescale methods that are commonly applied across neuroimaging modalities (fMRI, EEG, ECoG, MEG). The time-domain (TD) linear model estimated with linear least squares (Kaneoke et al., 2012; Meisel et al., 2017; Huang et al., 2018; Lurie et al., 2024; Shinn et al., 2023; Shafiei et al., 2020; Spitmaan et al., 2020; Trepka et al., 2024), and the autocorrelation-domain (AD) nonlinear model estimated with nonlinear least squares (Rossi-Pool et al., 2021; Cirillo et al., 2018; Ito et al., 2020; Runyan et al., 2017; Zeraati et al., 2022; Nougaret et al., 2021; Wasmuht et al., 2018; Müller et al., 2020; Maisson et al., 2021; Li and Wang, 2022; Shafiei et al., 2020). Both belong to the subclass of M-estimators defined by minimizing a sample squared error loss function. This approach seeks the best approximation in terms of squared error and does not require full specification of the data-generating distribution, relying on method of moments estimation.

Throughout, a superscript star (e.g. *τ*^∗^) denotes the pseudo-true parameter corresponding to the best fitting parameter in a possibly misspecified model. If correctly specified, *τ*^∗^ is the true timescale; otherwise, it is an approximation. Acknowledging that the data-generating process and fitted model will likely be different in practice, we introduce standard errors that account for this mismatch, enabling valid inference despite misspecification of the autocorrelation decay. Under the above assumptions and relevant moment conditions, the M-estimators are consistent for their respective pseudo-true parameters and asymptotically normal (see Estimator Properties). Further, they are invariant under the reparameterization needed to define timescales, with asymptotic normality preserved by the delta method, ensuring valid inference for transformed parameters.

### 2.2 Timescale Definitions

#### 2.2.1 Time-Domain Linear Model

A first-order autoregressive model (AR1) provides a linear approximation of the timescale. The AR1 model:

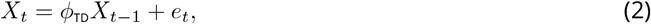

defines the process as a linear regression between *X*_*t*_ and *X*_*t*−1_ with *iid* innovations *e*_*t*_ with variance 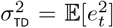. In the autocorrelation domain, it implies that the theoretical ACF decays exponentially at a rate determined by *ϕ*_TD_, such that 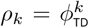 (Hansen, 2022, Chapter 14.22). For a stationary process with |*ϕ*_TD_ | < 1, the exponential decay rate can be directly obtained from *ϕ*_TD_, with a timescale *τ*_TD_ equal to the lag at which the AR1-projected ACF reaches 1/*e ≈* 0.37, that is, 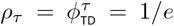, resulting in *τ*_TD_ = *g*(*ϕ*_TD_) = −1/log (|*ϕ*_TD_|). Here, the timescale *τ*_TD_ is expressed as a nonlinear function of *ϕ*_TD_, denoted by *g*(*ϕ*_TD_). This defines *τ*_TD_ to be a real number even though the ACF only includes integer indices. The absolute value allows for *ϕ*_TD_ < 0 and is introduced as a modeling assumption to ensure the timescale captures the monotonic decay rate of autocorrelations, consistent with the approach of Murray et al. (2014).

The parameter 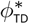 is the value that minimizes the expected squared error function *S*(*ϕ*_TD_):

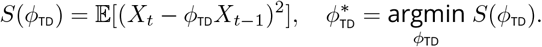

*S*(*ϕ*_TD_) is minimized by taking its derivative with respect to *ϕ*_TD_, setting it to zero, and solving for 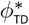 :

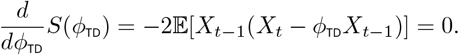

Differentiating the quadratic function yields a linear equation in *ϕ*_TD_, and solving this results in a closed-form expression for the optimal 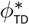 . Therefore, 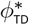 is defined by *linear projection* and the timescale parameter 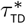 by a change of variable:

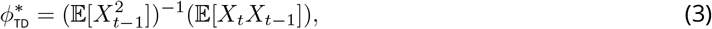

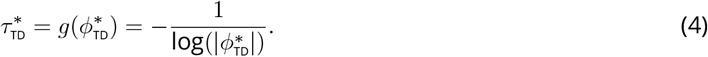

Thus, the timescale represents the *e*-folding time derived from the best-fitting AR1 model under squared error loss. Since *X*_*t*_ is stationary with finite variance, the parameters 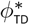 and 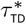 defined by projection are unique; in fact, any approximating AR1 model is identifiable if 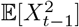 is nonzero and finite (Hansen, 2022, Theorem 14.28). Importantly, the observed process *X*_*t*_ need not follow the AR1 model from equation (2) with *iid* innovations *e*_*t*_. This allows for projection errors, i.e., residuals from linear projection onto the working model:

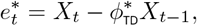

which may exhibit unequal variance and autocorrelation. This applies to any stationary and mixing process, even if the true data-generating process is not AR1, so the resulting fit is an AR1 projection.

#### 2.2.2 Autocorrelation-Domain Nonlinear Model

Alternatively, timescales can be defined in the autocorrelation domain by an exponential decay function, as introduced by Murray et al. (2014). We formalize this by projecting the *K*-lagged sequence of autocorrelations *ρ*_1_, …, *ρ*_*K*_ from (1) onto the exponential approximation 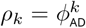 with decay parameter *ϕ*_AD_, minimizing the sum of squared errors averaged over fixed *K*:

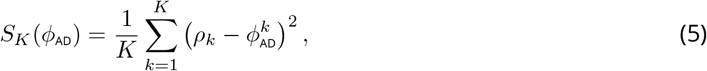

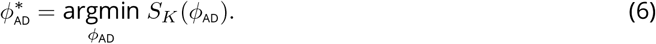

Unlike the Time-Domain Linear Model, which effectively projects using only a single (*K* = 1) lag of the ACF, this definition captures exponential decay across a sequence of multiple (*K*) lags. Consequently, the relationship between the pseudo-true parameters depends on the process specification:

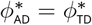 if correctly specified (AR1), whereas 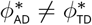 if misspecified. Crucially, because *ρ*_*k*_ and 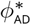 are deterministic population parameters rather than random variables, the projection residuals 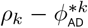 are fixed quantities distinct from the stochastic errors in the time domain. Under correct specification, these residuals vanish to zero; under misspecification, they quantify the systematic discrepancy between the complex true autocorrelation structure (e.g., multi-exponential or damped periodic) and the projected single-parameter exponential decay.

*S*_*K*_(*ϕ*_AD_) is minimized by taking its derivative with respect to *ϕ*_AD_, setting it to zero, and solving for *ϕ*_AD_ :

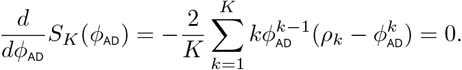

However, this first-order condition sums nonlinear terms across *K* lags, resulting in a high-order equation with no general closed-form solution. As a result, the pseudo-true parameter 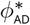 is defined implicitly as the solution to this nonlinear equation and computed using numerical optimization. As in the time-domain case, the corresponding timescale is defined as the *e*-folding time, expressed by the change of variable:

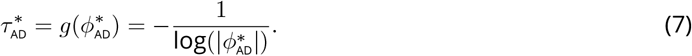

### 2.3 Timescale Estimation

#### 2.3.1 Time-Domain Linear Estimator

Given observations *x*_1_, …, *x*_*T*_ are demeaned, the linear least squares estimator of the Time-Domain Linear Model is obtained by replacing the expectations in equation (3) using method of moments. It has the following closed-form expression:

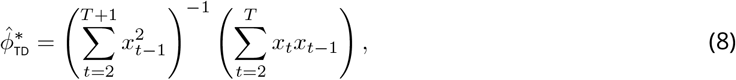

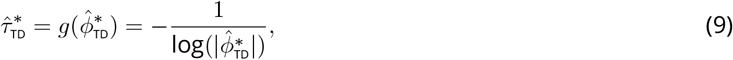

where (8) and (9) are the sample versions of the population parameters from equations (3) and (4), respectively (Hansen, 2022, Chapter 14.3).

#### 2.3.2 Autocorrelation-Domain Nonlinear Estimator

The nonlinear least squares estimator of the Autocorrelation-Domain Nonlinear Model is fit to the ACF, so the time series needs to be first transformed into the autocorrelation domain. For a finite-length and centered time series, the population ACF from equation (1) is estimated by:

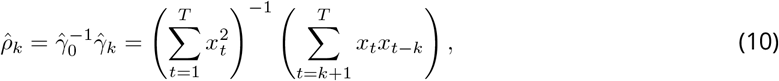

where 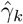 is the sample covariance at lag *k* and 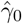 is the sample variance. Under the mixing assumption, the population ACF (1) approaches zero as lag *k* increases. However, sampling variability may yield non-zero autocorrelations even when true values are zero. To mitigate this, the sample ACF estimator (10) imposes a bias towards zero by scaling the autocovariances (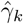, calculated using *T* − *k* terms) by the total sample variance (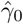, calculated using all *T* timepoints). Marriott and Pope (1954) demonstrated that this estimator minimizes bias and variance compared to other standard approaches in finite samples.

The exponential decay parameter 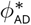 that minimizes the cost function *S*_*K*_(*ϕ*_AD_) in equation (5) is estimated by minimizing the sample analog 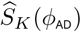:

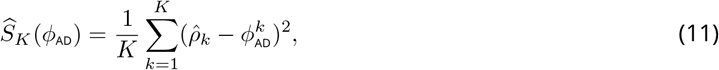

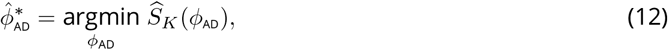

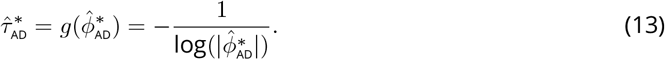

Here, (12) and (13) are the sample versions of the population parameters from equations (6) and (7), respectively. In this paper we use the Levenberg-Marquardt algorithm to iteratively update the estimate of 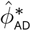 until convergence (i.e., when the step size goes below a 10^−6^ tolerance) (Levenberg, 1944).

### 2.4 Estimator Properties

In this section, we describe the large-sample properties of both the Time-Domain Linear Model and Autocorrelation-Domain Nonlinear Model, focusing on the consistency and asymptotic distribution of their respective estimators. Under general conditions — when the time-domain method is applied to a process that is not AR1, or the autocorrelation-domain method is applied to a decay process that is not exponential — we demonstrate that the asymptotic distribution is Gaussian, with a limiting variance that can be consistently estimated. Specific expressions for this variance and its estimator are provided after in sections Standard Error of the Estimators and Estimation of Standard Errors. This allows for the construction of hypothesis tests and confidence intervals across timescale maps of the brain.

#### 2.4.1 Time-Domain Method

Following the description in Hansen (2022, Theorem 14.29), the ergodic theorem shows that mixing (which implies ergodicity) is sufficient for consistent estimation. Since *X*_*t*_ is stationary and ergodic, so too are *X*_*t*_*X*_*t*−1_ and 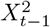, and therefore:

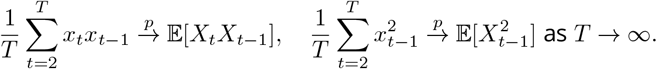

Applying the continuous mapping theorem yields:

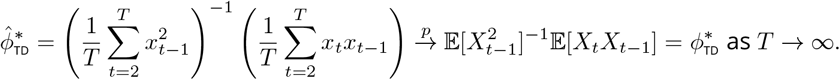

This shows that the coefficients of the Time-Domain Linear Model can be consistently estimated by least squares, for any stationary and mixing process with parameters defined by projection in equation (3).

Following Hansen (2022, Theorem 14.33), the asymptotic distribution under general dependence states that the behavior of the estimators of 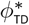 and 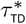 can be approximated using a central limit theorem for correlated observations:

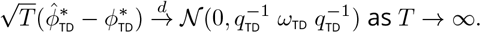

The estimator converges to a Gaussian distribution with a sandwich-form variance with components *q*_TD_ and *ω*_TD_ defined below in equation (14). And by the delta method we obtain the asymptotic distribution for the timescale given the transformation from equation (4):

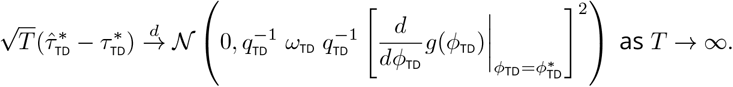

#### 2.4.2 Autocorrelation-Domain Method

To show consistent estimation, the sample autocorrelation 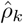 (10) is a ratio of sample moments in the time domain, and under the ergodic and continuous mapping theorems (Hansen, 2022, Theorem 14.9):

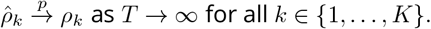

Consequently, the objective function (11) is itself a sample average, such that for any fixed *K* and *ϕ*_AD_ :

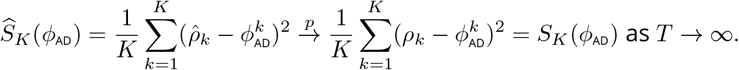

Further, if the minimizer 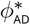 is unique, 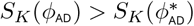 for all 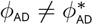 (Hansen, 2022, Theorem 22.1), then the sample minimizer from (12) converges to the population minimum:

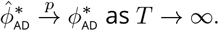

This shows that the parameters of the Autocorrelation-Domain Nonlinear Model can be consistently estimated by least squares. For this theoretical convergence to hold in practice, we assume that the numerical tolerance of the Levenberg-Marquardt algorithm approaches zero for large *T* .

With the additional assumption that the objective function (5) is Lipschitz-continuous for *ϕ*_AD_ near 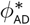, following Hansen (2022, Theorem 23.2), we can approximate the behavior of the estimators of 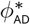 and 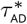 adapting the central limit theorem for nonlinear least squares. Since the estimator 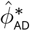 satisfies the first-order condition 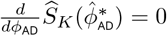, a mean-value expansion of the first derivative (score function) around the population parameter 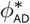 yields the linear approximation:

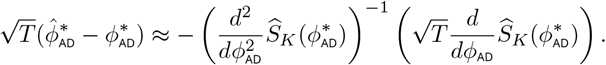

By the ergodic theorem, the sample second derivative (the first term) converges in probability to the deterministic curvature constant *q*_AD_ . In addition, the central limit theorem for mixing processes (Hansen, 2022, Theorem 14.33) implies that the scaled score (the second term) converges in distribution to a Gaussian with variance *ω*_AD_ . Combining these results yields the limiting distribution:

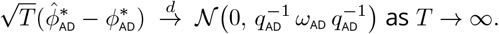

Thus, the autocorrelation-domain estimator exhibits Gaussian asymptotic behavior governed by a sandwich variance composed of *q*_AD_ and *ω*_AD_, which are defined below in equation (16). Finally, the asymptotic distribution of the timescale 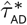 follows from the delta method, accounting for the nonlinear transformation in equation (7):

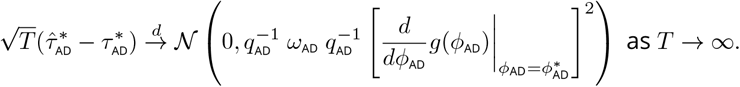

### 2.5 Standard Error of the Estimators

#### 2.5.1 Time-Domain Standard Error

Having established in Estimator Properties that the estimators are asymptotically normal, we now derive the specific variance components required for inference. When the data-generating process deviates from AR1 such that the errors are dependent and/or heteroskedastic, the usual (naive) standard errors will be biased. As a result, confidence intervals or hypothesis tests that rely on them are invalid. To correct for this, the Newey and West (1987) (NW) expression takes a sandwich form and explicitly accounts for misspecification by summing the covariance structure of the errors, ensuring that the resulting standard errors are asymptotically valid (Hansen, 2022, Theorem 14.32).

Given that *X*_*t*_ is stationary and mixing, so too are the errors from equation (2) since these properties are preserved by finite transformations (Hansen, 2022, Theorem 14.2 and Theorem 14.12). Consequently, the autocovariances of the errors vanish as the time lag increases (see Assumptions). Further, because the timescale estimator 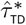 is given by the nonlinear function 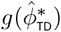, its theoretical standard error can be approximated by the delta method, which uses the derivative with respect to *ϕ*_TD_ evaluated at the pseudo-true parameter value 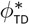 :

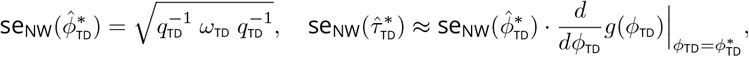

Where

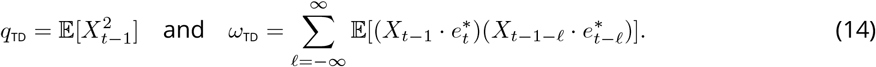

These moments, *q*_TD_ and *ω*_TD_, are stationary and therefore do not depend on time index *t*. The covariance terms in *ω*_TD_ capture deviations in the error structure from the standard *iid* case. For the special case of correct specification, when *X*_*t*_ is a true AR1 process, the standard error of 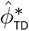 reduces to the usual formula:

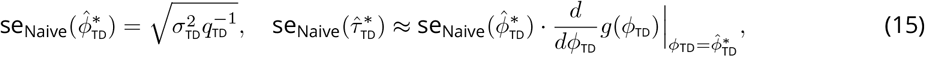

where 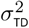 is the time-domain error variance.

#### 2.5.2 Autocorrelation-Domain Standard Error

Equation (5) models autocorrelation decay using a parametric exponential function. The standard errors proposed below account for potential misspecification of this decay (e.g., if the true process is not strictly exponential) and are asymptotically valid for any stationary and mixing process.

Following Hansen (2022, Chapter 22.8 and Chapter 23.5), the theoretical standard error of the estimator 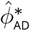 takes a Newey and West (1987) (NW) sandwich form that reflects both the curvature of the squared loss function and the time-domain covariance of the scores. The standard error for the timescale 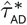 is then approximated via the delta method:

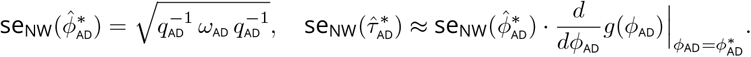

The curvature term *q*_AD_ is the sum of the squared first derivatives of the regression function 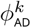 with respect to *ϕ*_AD_, evaluated at the minimizer 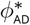 . The variance term *ω*_AD_ is the long-run variance of the normalized score process *u*_*t*_ (see Appendix for the derivation). This process is a weighted sum of the time-domain score contributions across all *K* lags, where the weights are given by the regression function derivative:

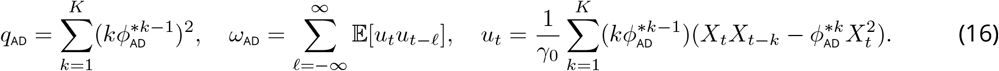

Defining *ω*_AD_ in terms of the score process *u*_*t*_ maps the joint dependence structure of the autocorrelation sequence *ρ*_1_, …, *ρ*_*K*_ onto a scalar process. Of note, *u*_*t*_ is defined in the time domain since *ρ*_*k*_ = *γ*_*k*_/*γ*_0_ is a function of moments of *X*_*t*_. This linearized representation ensures that the long-run variance *ω*_AD_ captures the relevant dependencies, including both the correlations within and between lags. For a stationary process with 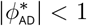, the weights 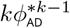 in the score process decrease rapidly with lag *k*, meaning the asymptotic variance is largely determined by the variability of the lower-order sample autocorrelations. Under the special case of correct AR1 specification, higher-order lags provide no additional information as they are deterministic functions of the first lag *ρ*_1_. In this case, the autocorrelation-domain standard error is asymptotically equivalent to the time-domain standard error in (15), replacing 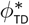 with 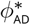 .

### 2.6 Estimation of Standard Errors

#### 2.6.1 Time-Domain Standard Error Estimator

The sample standard error estimator takes the form:

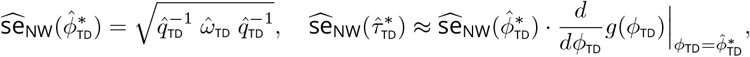

Where

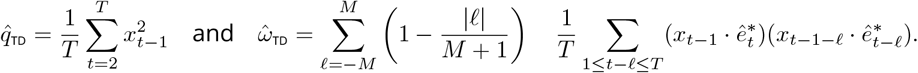

This estimator calculates a weighted sum of the regression scores 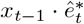, where 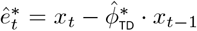. The true *ω*_TD_ is approximated by 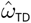 by taking a finite sum of the regression score covariances up to lag *M*, where *M* is the lag-truncation (or bandwidth). The weights used in the sum decrease linearly with lag *𝓁*, following a Bartlett kernel (Newey and West, 1987). This kernel not only ensures the standard errors remain non-negative but also regularizes 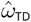 to change smoothly with *M* (Hansen, 2022, Chapter 14.35).

For comparison we also include the naive estimator which simplifies under correct specification:

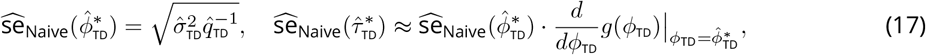

where 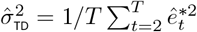 is an estimate of the time-domain error variance.

#### 2.6.2 Autocorrelation-Domain Standard Error Estimator

The sample standard error estimator takes the form:

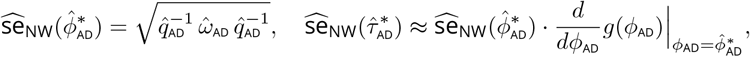

where the curvature 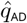 is estimated by the sum of the squared first derivatives evaluated at 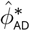, and the long-run variance *ω*_AD_ is obtained by applying a Newey–West estimator to the sample normalized score process *û*_*t*_:

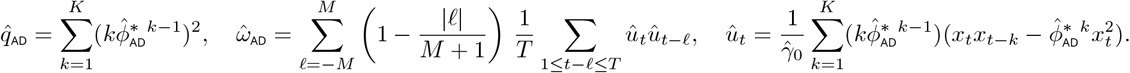

The estimator 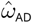 sums sample autocovariances of *û*_*t*_ up to lag *M* using a Bartlett kernel. This explicitly accounts for serial dependence in *x*_*t*_ that is not captured by the fitted autocorrelation model, ensuring robust inference under misspecification (e.g., non-exponential decay). In the case of correct specification, the equation simplifies to (17), replacing 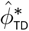 with 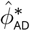 .

## 3 Simulations

### 3.1 Simulation Settings

Monte Carlo simulations with *B* = 10, 000 replications were used to evaluate time- and autocorrelation-domain methods. Time series realizations *x* = {*x*_1_, *x*_2_, …, *x*_*T*_} with *T* = 4800 were based on three data-generating processes, each characterized by a different autocorrelation structure: AR1, AR2, and autocorrelations derived from rfMRI data. All autocorrelation structures shared the same time-domain projection parameter 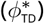 for comparable timescales. That is, there is always a 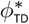 value that represents the time-domain AR1 model projection, even if the time series was generated by a more complex process. To define a feasible parameter range for simulation, we referred to the Human Connectome Project (HCP) dataset, where 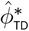 estimates ranged from +0.1 to +0.8. Accordingly, autocorrelation strength was varied using five positive 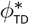 values (0.1 to 0.8) with corresponding timescales 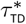 from equation (4). This design resulted in a total of 15 simulation settings (three data-generating models *×* five autocorrelation strengths). For each setting, estimator performance was assessed by relative root mean squared error (rRMSE):

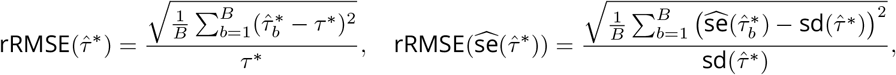

where 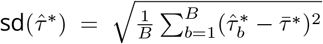 is the empirical standard deviation of the timescale estimator across *B* simulation replications, and 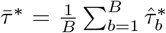 . The rRMSE captures both bias and variance of timescale estimates and their standard errors as a proportion of the true value, facilitating comparison across parameter ranges and simulation settings. Cutoffs of <10% rRMSE for timescales and <20% for standard errors are shown to indicate “good” estimation, following applied time series conventions (Jadon et al., 2024).

In the *AR1 setting*, the data-generating process matches the fitted model, with time series simulated from the AR1 model:

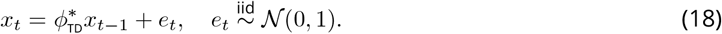

The *AR2 setting* misspecifies the order of the autoregression by introducing a mismatch between the AR2 data-generating process and AR1 fitting, with five pairs of AR2 coefficients selected so that the 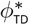 projection matched the above setting (see Figure 2 Panel A). The following model was used:

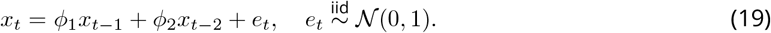

The *HCP setting* did not follow an autoregressive process, and instead used empirical ACFs from five brain regions of subject #100610 in the HCP dataset (see Dataset Description). These regions were selected to match the 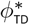 projections above. To simulate time series with the same autocorrelation structure as the empirical data, we sampled from a multivariate normal distribution 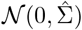, where 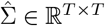 is the covariance matrix constructed from the sample ACFs. Under stationarity, 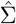 is Toeplitz, so its *k*^th^ off-diagonal equals the sample ACF at lag 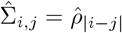 for indices *i, j* = *{*1, …, *T }*. A Cholesky decomposition of the covariance matrix 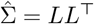, where *L* is a lower triangular matrix, was multiplied with Gaussian white noise:

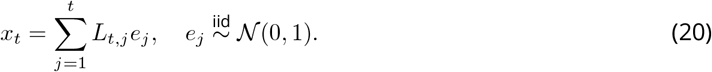

### 3.2 Simulation Results

#### 3.2.1 Results for Autoregressive Simulations

AR1 simulations (**Figure 1**) serve as a baseline for correct specification, where the data-generating process defined in (18) aligns with the fitted models (**Panel A**). **Panel B** demonstrates that while estimation variance increases with the true timescale, rRMSE remains below the conventional 10% threshold. Regarding uncertainty quantification (**Panels C-D**), the time-domain estimator **(Row 1)** shows minimal difference between Naive and Newey-West approaches because of correct specification. However, the autocorrelation-domain estimator **(Row 2)** exhibits a downward bias that is mitigated by the Newey-West correction. This bias likely occurs because fitting multiple noisy lags in the autocorrelation domain introduces additional sampling variability that the naive formula does not capture. Consistent with this, the autocorrelation-domain method produces point estimates and standard errors with higher overall variance than the time-domain method.

**Figure 1.**
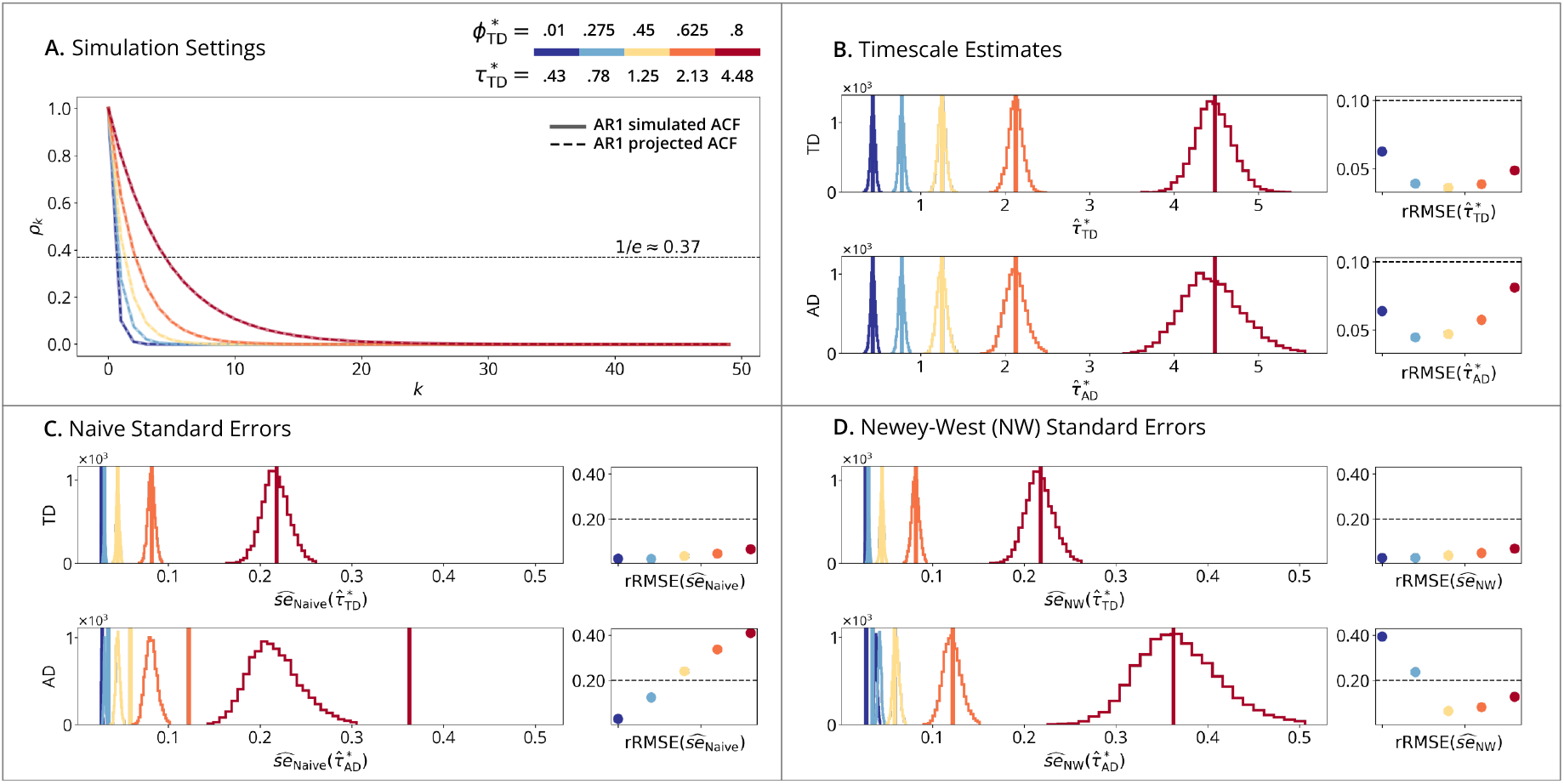
AR1 time-domain simulations. (**A**) Simulation Setting: solid lines show the simulated ACFs of the process defined in equation (18); dashed lines show the AR1-projected ACFs which are overlapping as both follow AR1. Horizontal line marks the timescale where the AR1-projected ACF reaches 1/*e ≈* 0.37. (**B**) Timescale Estimates: vertical lines show true timescales *τ*^∗^; histograms show estimates 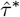 across *B* = 10, 000 replications; points show rRMSE versus a 10% error line. (**Row 1**): Time-Domain Linear Estimator. (**Row 2**): Autocorrelation-Domain Nonlinear Estimator. (**C**) Naive and (**D**) Newey-West Standard Errors: vertical lines show true standard errors 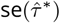, i.e., standard deviations of the sampling distributions from panel B; histograms show standard error estimates 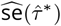 ; points show rRMSE versus a 20% error line. (**Row 1**): Time-Domain Standard Error Estimator. (**Row 2**): Autocorrelation-Domain Standard Error Estimator.

AR2 simulations (**Figure 2**) assess estimator performance under model misspecification, using a higher-order process defined in (19) that deviates from the fitted AR1 model (**Panel A**). **Panel B** shows that the two estimators converge to different timescales due to their distinct definitions. In terms of standard errors, **Panel C** shows a consistent underestimation by the naive estimator across both methods because of misspecification. **Panel D** demonstrates that this bias is largely mitigated by the Newey-West estimator.

**Figure 2.**
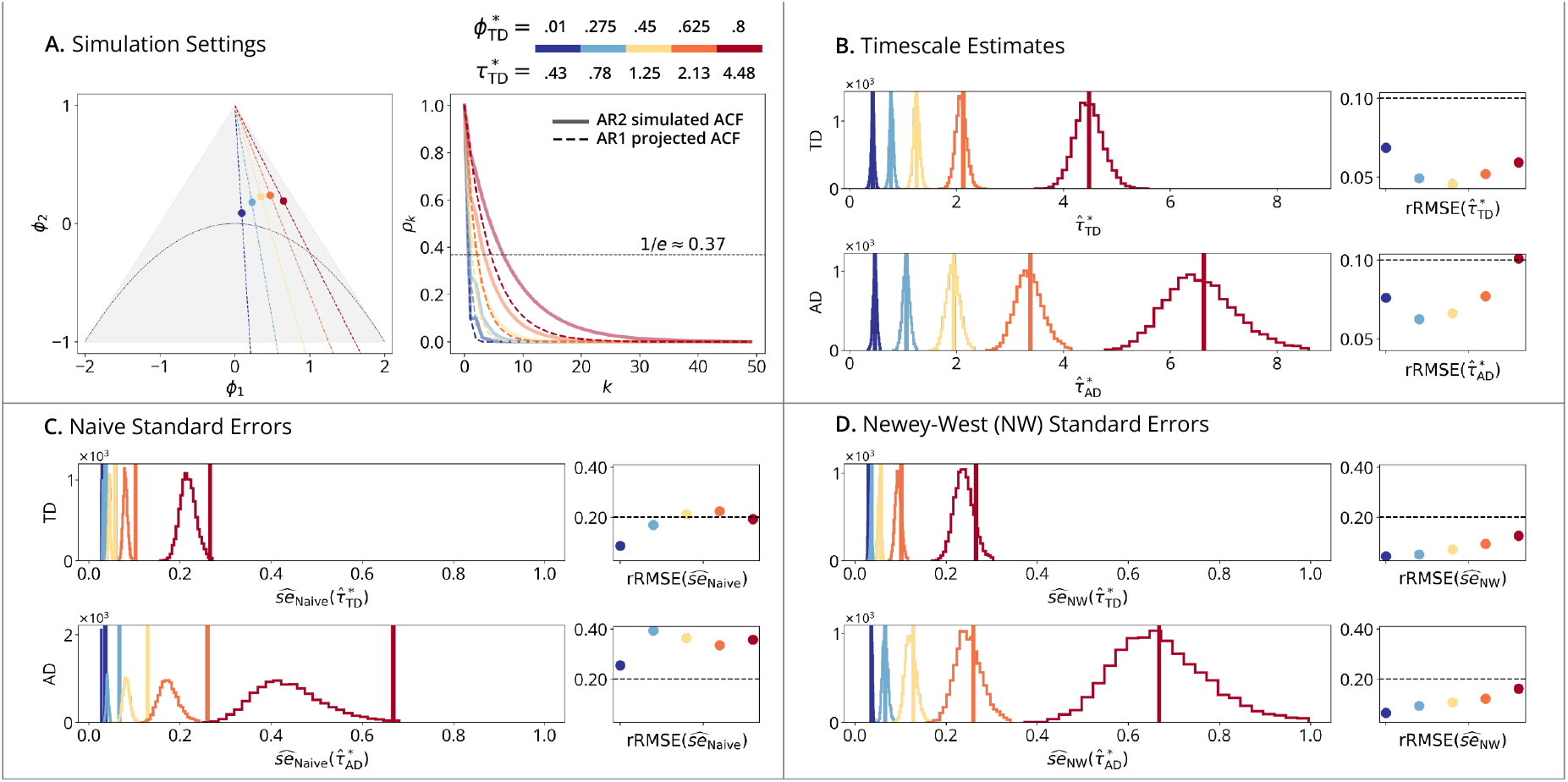
AR2 time-domain simulations. (**A**) Simulation Setting (**Left**): triangle shows the AR2 stationary region in the (*ϕ*_1_, *ϕ*_2_) plane, with the periodic/ape-riodic boundary at 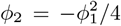 . Dashed lines within the triangle illustrate the preimage under the AR2-to-AR1 projection mapping; specifically, all (*ϕ*_1_, *ϕ*_2_) pairs on a given dashed line correspond to AR2 processes that project to the same AR1. Points show five selected AR2 (*ϕ*_1_, *ϕ*_2_) pairs, with their corresponding AR1 projections shown in the colorbar. (**Right**): solid lines plot the simulated ACFs from the process defined in equation (19); dashed lines show the ACFs based on AR1-projected models, which are not overlapping because the simulated AR2 differs from its AR1 fit. (**B**) Timescale Estimates: vertical lines show true timescales *τ*^∗^; histograms show estimates 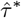 across *B* = 10, 000 replications; points show rRMSE versus a 10% error line. (**Row 1**): Time-Domain Linear Estimator. (**Row 2**): Autocorrelation-Domain Nonlinear Estimator. (**C**) Naive and (**D**) Newey-West Standard Errors: vertical lines show true standard errors 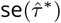, i.e., standard deviations of the sampling distributions from panel B; histograms show standard error estimates 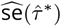 ; points show rRMSE versus a 20% error line. (**Row 1**): Time-Domain Standard Error Estimator. (**Row 2**): Autocorrelation-Domain Standard Error Estimator.

#### 3.2.2 Results for Realistic rfMRI Simulations

Realistic rfMRI simulations (**Figure 3**) evaluate estimator performance using the data-generating process defined in (20) with realistic autocorrelation derived from five distinct brain regions. Similar to the AR2 setting, this analysis examines the impact of model misspecification, as the empirical ACFs differ from the projected AR1 fits (**Panel A**). **Panel B** confirms that time- and autocorrelation-domain estimators capture distinct timescale values due to their respective definitions. **Panels C-D** further demonstrate that naive standard errors are systematically underestimated, a bias that is substantially mitigated by the Newey-West correction.

**Figure 3.**
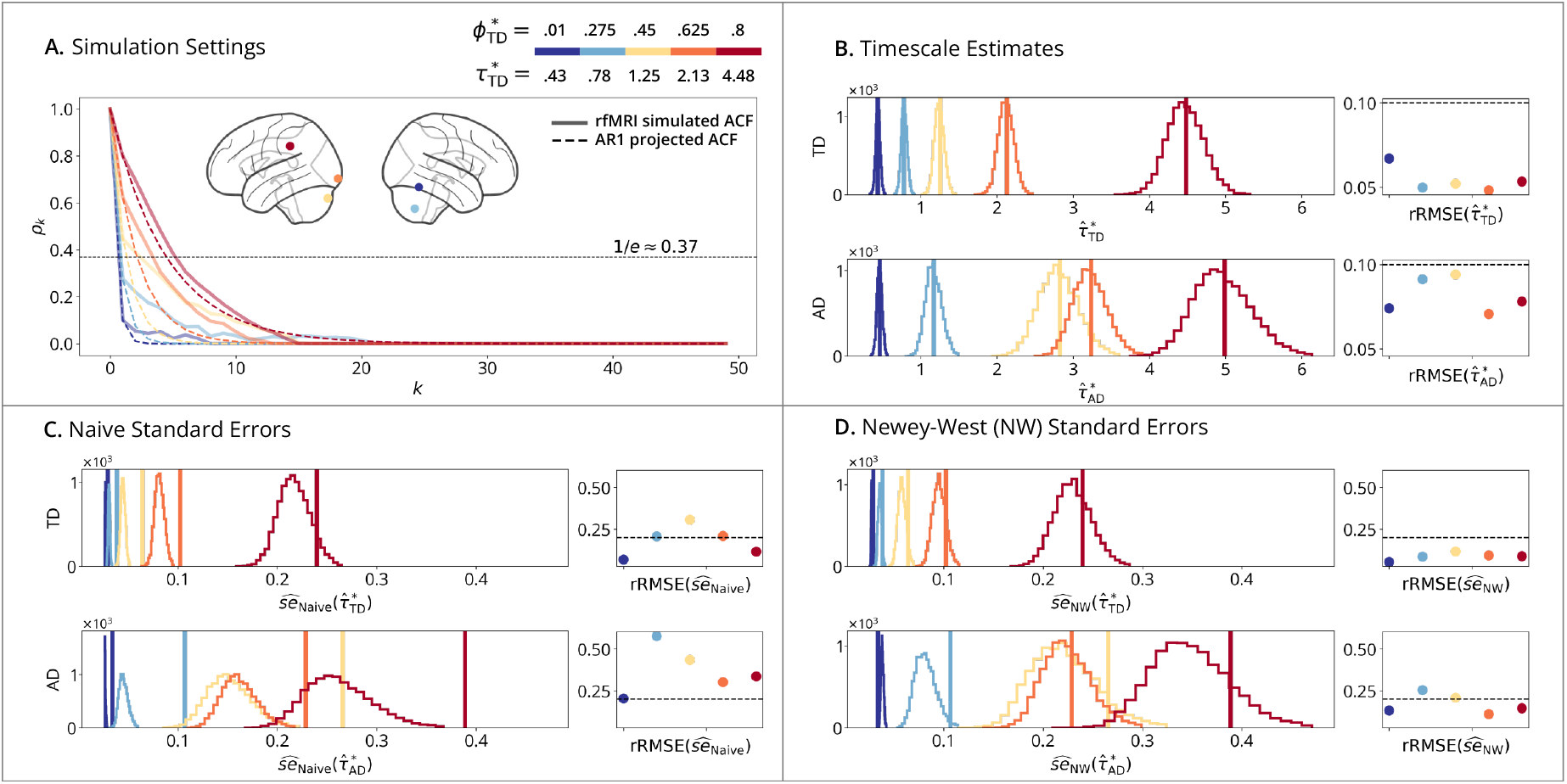
Realistic rfMRI time-domain simulations. (**A**) Simulation Setting: solid lines show estimated ACFs from five selected brain regions of HCP subject #100610 used to generate the process define in equation (20); dashed lines show the AR1-projected ACFs, which are not overlapping because the empirically estimated ACF differs from its AR1 fit. (**B**) Timescale Estimates: vertical lines show true timescales *τ*^∗^; histograms show estimates 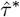 across *B* = 10, 000 replications; points show rRMSE versus a 10% error line. (**Row 1**): Time-Domain Linear Estimator. (**Row 2**): Autocorrelation-Domain Nonlinear Estimator. (**C**) Naive and (**D**) Newey-West Standard Errors: vertical lines show true standard errors 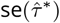, i.e., standard deviations of the sampling distributions from panel B; histograms show standard error estimates 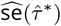 ; points show rRMSE versus a 20% error line. (**Row 1**): Time-Domain Standard Error Estimator. (**Row 2**): Autocorrelation-Domain Standard Error Estimator.

## 4 Data Analysis

### 4.1 Dataset Description

Resting fMRI (rfMRI) scans were provided by the Human Connectome Project (HCP), WU-Minn Consortium (led by principal investigators David Van Essen and Kamil Ugurbil; 1U54MH091657) funded by the 16 NIH Institutes and Centers supporting the NIH Blueprint for Neuroscience Research, and by the McDonnell Center for Systems Neuroscience at Washington University (Van Essen et al., 2013). Informed consent was obtained from all participants. Two subsets of the dataset were used: one for methods development and defining realistic simulation parameters (see Simulations), and the other for estimating high-resolution timescale maps of the cortex.

The *methods development subset* included 10 subjects (#100004 - #101410) scanned with a 3T gradient-echo EPI sequence (TR=720ms, slice thickness=2mm). Each subject completed four 15-minute runs (4800 timepoints total), preprocessed with standard steps including motion regression and artifact removal with ICA-FIX (see Glasser et al. (2013); Salimi-Khorshidi et al. (2014) for details). To further mitigate systemic low-frequency physiological oscillations driven by respiration and CO_2_ variation, we applied RAPIDtide (Frederick, 2025). This step accounts for the regionally varying lags of these global signals, caused by differences in blood arrival times, which can otherwise mimic or inflate regionally specific autocorrelation. The four runs were concatenated into a single continuous time series, under the assumption of weak stationarity (constant mean and variance) across denoised runs. The resulting dataset dimensions were {10 subjects, 4800 timepoints, 300 regions}. The *timescale mapping subset* included 180 subjects scanned with a 7T gradient-echo EPI sequence (TR=1000ms, slice thickness=1.6mm) over four 16-minute runs (3600 timepoints total), using the same preprocessing and concatenation steps. Functional data were analyzed on the cortical surface downsampled to 2mm spatial resolution, yielding a dataset with the dimensions {180 subjects, 3600 timepoints, 64984 vertices}. The time- and autocorrelation-domain methods were fit to each vertex independently, a mass-univariate analysis approach that resulted in subject-level spatial maps of temporal timescale estimates and their standard errors.

#### 4.1.1 Group-level Analysis

Group-level maps combined individual timescales and standard errors, accounting for within- and between-subject variability. For the group-level analysis, we used the *timescale mapping subset* of *N* = 180 subjects *T* = 3600 concatenated timepoints, distinct from the *methods development subset* of *N* = 10 subjects described above. While remaining within the mass-univariate framework, for simplicity, we express the group timescale at a single cortical vertex:

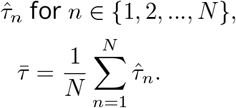

The group-level standard error for the timescale is given by the law of total variance:

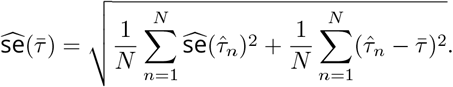

Here, the first term under the square root is the within-individual variance and the second term is the between-individual variance.

For visualization, brain-wide t-statistic maps tested whether timescales exceeded 0.5 seconds under the null hypothesis *H*_0_ : *τ* ≤ 0.5, by comparing the empirical group mean 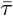 to the null value 0.5:

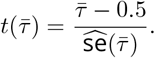

Similarly, relative standard error (RSE) maps visualize the spatial precision of timescale estimates to show the reciprocal of the unadjusted (zero-valued null) t-ratio:

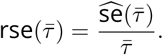

### 4.2 Data Analysis Results

#### 4.2.1 Results for rfMRI Timescale Maps

Subject-level maps (**Figure 4**) were generated by mass-univariate fitting of time- and autocorrelation-domain estimators to cortical surface data at the individual vertex level (64984 vertices). **Panel A** presents the spatial distribution of timescale estimates, which show that autocorrelation-domain estimates tend to yield larger timescales than the time-domain estimator, consistent with simulation results. **Panel B** shows the corresponding maps of standard errors, which are aligned with the timescale maps, consistent with the simulation finding that larger timescales are associated with greater sampling variability. **Panel C** depicts the relative ratio of timescale estimates to their standard errors (i.e., t-statistics), testing whether the timescales significantly exceed a half second (*H*_0_ : *τ* ≤ 0.5). Despite larger standard errors for higher timescales, these regions still exhibit higher t-statistics. **Panel D** shows low RSEs across much of the brain indicating high estimation reliability.

**Figure 4.**
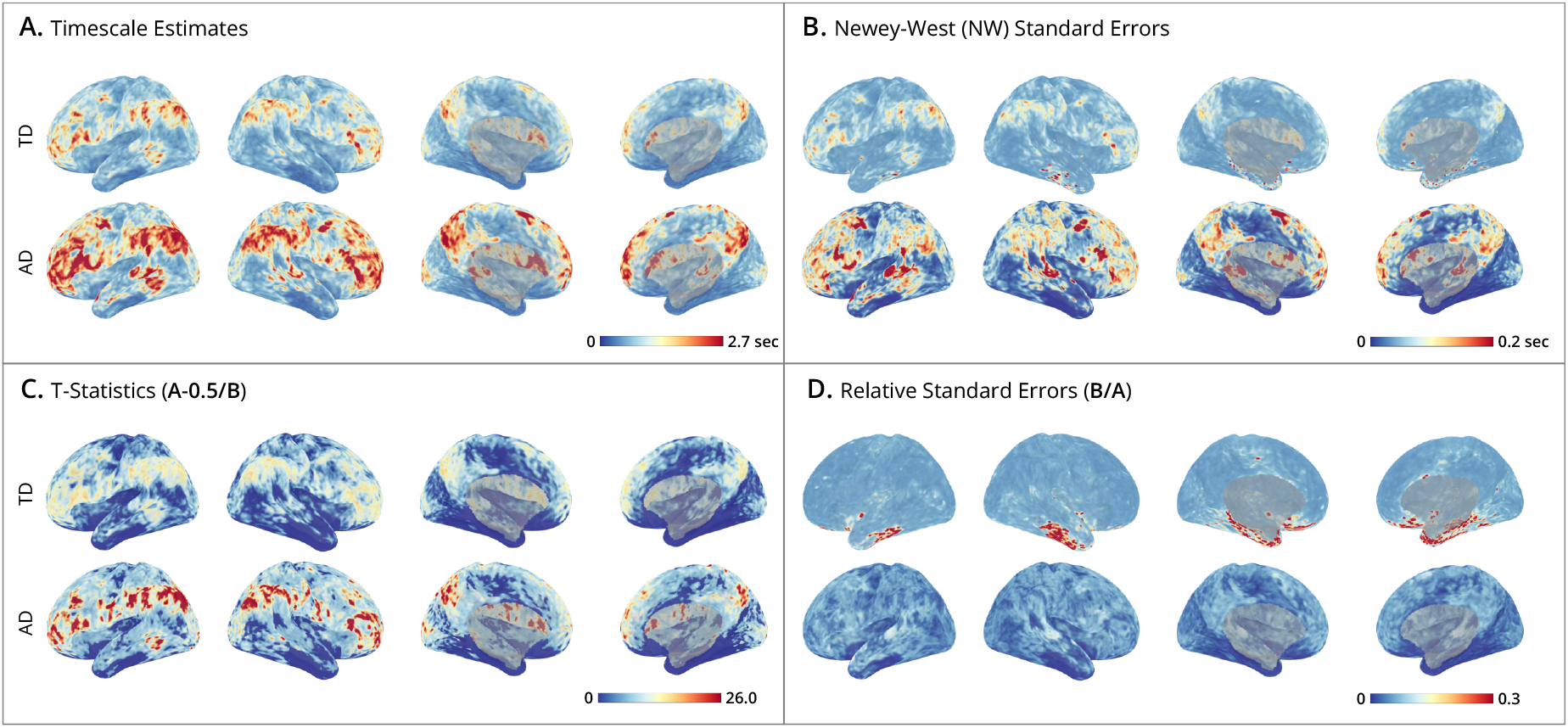
Human Connectome Project subject-level timescale maps. (**A-D**) Cortical surface maps from HCP subject #100610. Displays show lateral-left, lateral-right, medial-left, and medial-right views, plotted using the *StatBrainz* package (Davenport, 2025). The upper bounds on the colorbars are set for each panel at the 99^th^ percentile of cortical map values. For each panel, the top row shows results of the time-domain method, and the bottom row the autocorrelation-domain method. (**A**) Timescale estimates: maps display the timescales (in seconds) estimated at each vertex. (**B**) Newey-West standard errors: shows the spatial distribution of standard errors, where smaller values indicate greater estimation precision. (**C**) T-statistics: unthresholded and uncorrected t-ratios testing where timescales exceed 0.5 seconds. (**D**) Relative Standard Errors (RSEs): relative reliability of estimates, where low RSE (near zero) indicates high precision with small uncertainty.

Group-level timescale maps (**Figure 5**) were generated by combining individual estimates at the vertex level to account for both within- and between-individual variability, providing an aggregate view of timescale distributions across subjects. The smooth appearance of these timescale maps reflects the high spatial resolution of the vertex-level analysis and the averaging across subjects. **Panel A** shows that the average timescale maps for both time- and autocorrelation-domain estimators are smoother than the individual maps, displaying a well-organized spatial pattern across the cortex; notably autocorrelation-domain estimates are generally larger than time-domain estimates. **Panel B** presents the standard error maps, which combine variances from within-subject Newey-West estimates and between-subject timescale estimates. As expected from the simulation results, the standard errors are larger in the autocorrelation domain than in the time domain. **Panel C** depicts t-statistics testing whether timescales exceed half a second, while **Panel D** plots relative standard errors (RSEs), providing complementary views of estimation reliability. Interpreting these maps together clarifies how signal strength interacts with uncertainty: although standard errors theoretically increase with the true timescale (as shown in sections 2.5 and 3), these empirical results demonstrate that the estimated timescale magnitude grows faster than its associated error. Consequently, regions with longer timescales actually exhibit higher signal-to-noise ratios (larger t-statistics). In contrast, regions with shorter timescales, such as the limbic network (comprising orbitalfrontal cortex and anterior temporal cortex), exhibit greater relative uncertainty and lower signal-to-noise ratios (Yeo et al., 2011; Du et al., 2024).

**Figure 5.**
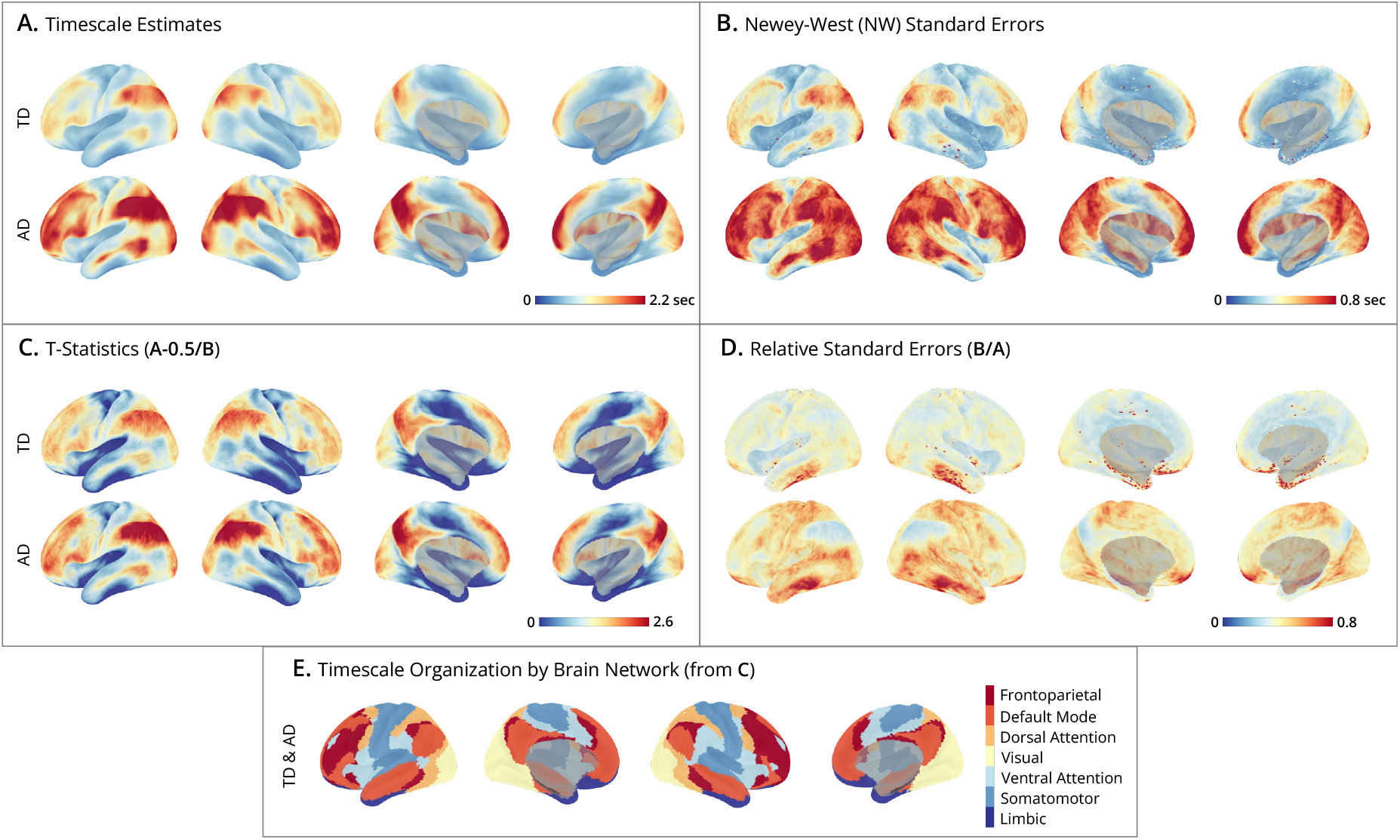
Human Connectome Project group-level timescale maps. (**A-D**) Cortical surface maps from *N* = 180 HCP subjects. Displays show lateral-left, lateral-right, medial-left, and medial-right views, plotted using the *StatBrainz* package (Davenport, 2025). The upper bounds on the colorbars are set for each panel at the 99^th^ percentile of cortical map values. For each panel, the top row shows results of the time-domain method, and the bottom row the autocorrelation-domain method. (**A**) Timescale estimates: maps display the group-level timescales (in seconds) at each vertex, averaged over subjects. (**B**) Newey-West standard errors: group-level spatial distribution of estimates, accounting for within- and between-subject variability. Smaller values indicate greater precision. (**C**) T-statistics: unthresholded and uncorrected t-ratios testing where group-level timescales exceed 0.5 seconds. (**D**) Relative standard errors (RSEs): relative reliability of estimates, where low RSE (near zero) indicates high precision with small uncertainty across subjects. (**E**) Timescale organization by brain network: maps display brain networks from the Yeo 7 Network Atlas, ordered by the network-averaged t-statistics (from panel C). This ordering is the same for time- and autocorrelation-domain methods, and highlights the hierarchical organization of timescales, progressing from sensory networks (e.g., somatomotor and limbic in blues) to association networks (e.g., frontoparietal and default mode in reds).

**Panel E** highlights the spatial organization of timescales into networks by mapping the t-statistic at each vertex to one of seven networks from the Yeo et al. (2011) atlas. The ordering {limbic, somatomotor, ventral attention, visual, dorsal attention, default, frontoparietal} aligns with the sensory-to-association axis of brain organization and is consistent with an extensive literature on the hierarchical organization of timescales (Murray et al., 2014; Stephens et al., 2013; Raut et al., 2020; Gao et al., 2020; Hasson et al., 2008). Sensory networks (limbic and somatomotor) have short timescales, followed by attentional networks (ventral and dorsal attention), while higher-order association networks (default mode and frontoparietal) show the longest timescales. However, interpreting these network differences requires caution, particularly because regions in the limbic network are especially susceptible to measurement noise which can conflate timescale estimation (see Effects of Measurement Noise).

This empirical analysis highlights methodological considerations for estimating fMRI timescale maps. (i) Both time- and autocorrelation-domain methods produce similar maps, but diverge at extremes – the autocorrelation-domain estimator yields larger timescales. The corresponding standard errors in the autocorrelation domain are also larger, so the resulting t-ratio maps remain similar between the two methods, highlighting why point estimates can be misleading without considering standard errors. Likewise, estimates are on average more precise for the time-domain estimator, consistent with simulation results. (ii) In this mass-univariate analysis, the computational cost of the time-domain estimator is substantially lower than autocorrelation-domain estimator because of its simple analytical solution. (iii) The t-statistic maps organized by brain network exhibit a clear hierarchical timescale organization, reflecting how networks integrate and process information over time. And this pattern is consistent regardless of the estimation method. (iv) As a sensitivity check, we repeated the analyses without added RAPIDtide denoising step and found that it had only a small influence on local vertex-level timescale estimates, and it did not alter the global network-level organization (e.g., the relative ordering of networks by timescale). Taken together, these findings suggest that the time-domain method may be preferable for large-scale neuroimaging studies due to its computational efficiency and higher precision, while producing maps consistent with previously reported timescale hierarchies, and that are robust to two different denoising strategies.

## 5 Practical Considerations for fMRI Timescale Estimation

In this section, we provide recommendations for practitioners seeking to estimate fMRI timescale maps. We advocate for the Time-Domain Linear Estimator over the Autocorrelation-Domain Nonlinear Estimator, as it is more computationally efficient while maintaining robust performance in Simulations. However, a critical distinction must be drawn between the *statistical validity* of the estimator and the *interpretability* of the resulting timescale maps, which inevitably reflect the composite nature of the observed fMRI BOLD signals. The mixed metabolic and neuronal origins of the hemodynamic signal complicate mechanistic interpretations (Raut et al., 2020; He, 2011). Consequently, interpreting fMRI timescales as proxies for *intrinsic neural timescales* requires disentangling latent neural dynamics from confounding sources of variability. The following subsections detail how the hemodynamic response function, measurement noise, and periodicity specifically impact these interpretations.

### 5.1 Effect of the Hemodynamic Response Function

The fMRI BOLD signal is a composite measure reflecting underlying neural activity filtered through a complex vascular system. While neural populations exhibit the characteristic exponential decay in autocorrelation that defines the intrinsic neural timescale (Murray et al., 2014, Supplementary Mathematical Note), the hemodynamic response function (HRF) acts as a temporal low-pass filter on this latent neural activity. Forward modeling indicates that while this filtering inflates absolute timescale values, it generally preserves the relative spatial hierarchy, recovering the sensory-to-association axis observed empirically (Watanabe et al., 2019).

However, reliance on a canonical HRF may oversimplify the relationship between neural and vascular dynamics. Recent evidence demonstrates that HRF shape varies across both brain regions and individuals (Bailes et al., 2023; Rangaprakash et al., 2018). This variability introduces a potential confound where observed timescale differences may stem from vascular origins, such as regional heterogeneity in blood flow latency, rather than differences in neural processing. A method proposed by Wu et al. (2021) for HRF estimation and deconvolution offers a promising route to disentangle these effects from the resting-state BOLD signal. Nevertheless, a fundamental identifiability problem remains in that both a wider HRF and a longer neural timescale can produce nearly identical observed BOLD signals. Resolving this ambiguity requires moving beyond observational studies, and future work should leverage experimental manipulations and/or concurrent EEG-fMRI to tease apart vascular and neural timescales.

### 5.2 Effects of Measurement Noise

While the HRF tends to lengthen timescales, measurement noise typically biases estimates in the opposite direction. In the fMRI setting, the observed time series is rarely generated by a clean autoregressive process; rather, it is a sum of distinct stochastic processes capturing neural activity, hemody-namic filtering, and measurement noise. In such settings, regions with identical latent neural timescales can yield divergent observed fMRI timescales depending on their local signal-to-noise ratio. This raises an interpretational concern where the fitted timescale may be statistically well-estimated yet fail to correspond to the intended parameter of neural integration if measurement noise dominates.

To illustrate this limitation, consider the simulation setting where the latent neural process is generated as an AR1 with a fixed timescale 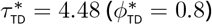 :

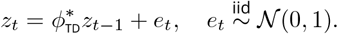

Observations are formed by adding independent measurement noise of increasing magnitude to vary observation quality while holding the latent timescale constant:

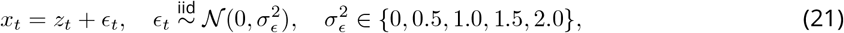

Under this construction, increasing *σ*^2^ accelerates the rate of autocorrelation decay (**Figure 6 Panel A**). This occurs because noise inflates the lag-0 variance without altering cross-lag covariances, shrinking the autocorrelation ratio in equation (1). Consequently, the time-domain estimator in **Panel B row 1** diverges from the true latent neural timescale, reflecting instead the *pseudo-true timescale* of the misspecified process (see Estimation and Inference under Misspecification). Importantly, while the Newey-West standard errors in **Panel B row 2** ensure that inference on this pseudo-true timescale remains statistically valid, its interpretation should reflect the mixture of neural integration and confounding measurement noise.

**Figure 6.**
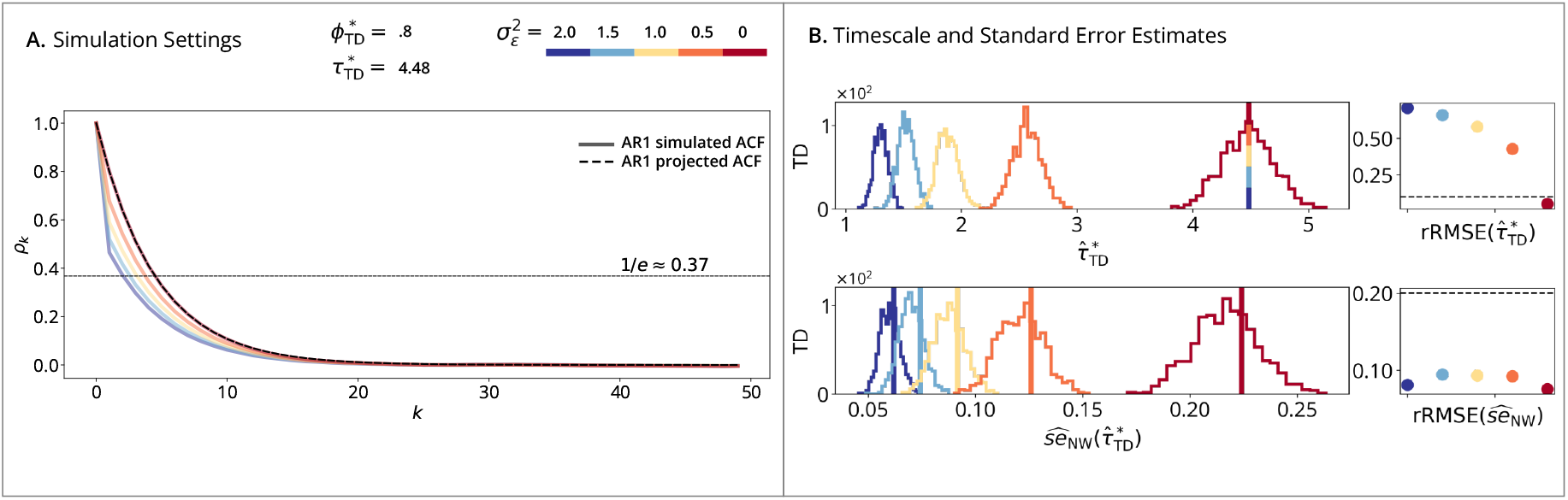
AR1 with added measurement noise simulations. (**A**) Simulation Setting: solid lines show the simulated ACFs of the AR1 + measurement noise process defined in equation (21) with a fixed timescale and increasing measurement noise variance 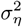 ; the dashed black line shows the AR1-projected ACF with no noise variance 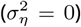 . Increasing noise (red-to-blue) increases the rate of autocorrelation decay even when the latent timescale is fixed. (**B**) Timescale and Standard Error Estimates: vertical lines show the true latent neural timescale (*τ*_TD_ = 4.48 for all settings); histograms show estimates across *B* = 10, 000 replications; points show rRMSE. **(Row 1)** Time-Domain Linear Estimator, where rRMSE is plotted versus a 10% error line. (**Row 2**) Time-Domain Standard Error Estimator (Newey-West), where rRMSE is plotted versus a 20% error line.

This dependency emphasizes that the fitted timescale is effectively a confounded measure of both neural dynamics and measurement noise and the resulting timescale should be interpreted as a pseudo-true timescale under model misspecification. Therefore in practice, effective denoising is a prerequisite for separating neural timescales from artifacts such as head motion and physiological noise. Regarding motion, the proposed Time-Domain Linear Estimator offers a practical advantage; while standard strategies often require censoring followed by interpolation (Goldberg et al., 2024), this estimator relies solely on adjacent timepoint pairs (*x*_*t*_, *x*_*t*+1_). This allows motion-contaminated frames to be excluded without interpolation. Finally, even with careful denoising, interpretational caution is warranted in regions susceptible to signal loss, such as the orbital frontal cortex and the anterior inferior medial temporal lobe (Yeo et al., 2011; Du et al., 2024).

### 5.3 Effects of Periodicity

In addition to the HRF and measurement noise, we consider the influence of periodic processes, since neural timescales are generally considered to originate from the brain’s aperiodic (non-rhythmic) activity. While monotonic autocorrelation decay is consistent with an AR1 process, latent neural activity may exhibit rhythmic dynamics (He et al., 2010; He, 2011). Furthermore, non-neural sources, such as respiration and CO_2_ variation, can introduce systemic low-frequency oscillations (Tong et al., 2019; Korponay et al., 2024). These dynamics manifest as periodic fluctuations in the ACF, creating a case of model misspecification. To validate the robustness of the time-domain estimator in this setting, we simulated AR2 processes that display oscillations, holding the projected timescale constant 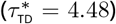 while increasing the oscillatory strength via the second-order coefficient 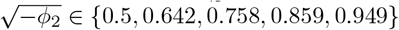 .

As shown in **Figure 7 Panel A**, we selected parameters along a preimage of constant timescale but increasing periodicity. Despite the mismatch between the data generating process (AR2) and the estimator (AR1), the histograms in **Panel B row 1** are accurately centered on the pseudo-true timescale. This demonstrates that the estimator recovers the dominant decay rate even amidst substantial oscillations. Notably, increasing the strength of oscillations decreases the variance of the sampling distributions; as the process becomes more rhythmic, the time-domain error variance shrinks. Consistent with this, the Newey-West standard errors (**Panel B row 2**) accurately track this reduction in sampling variability.

**Figure 7.**
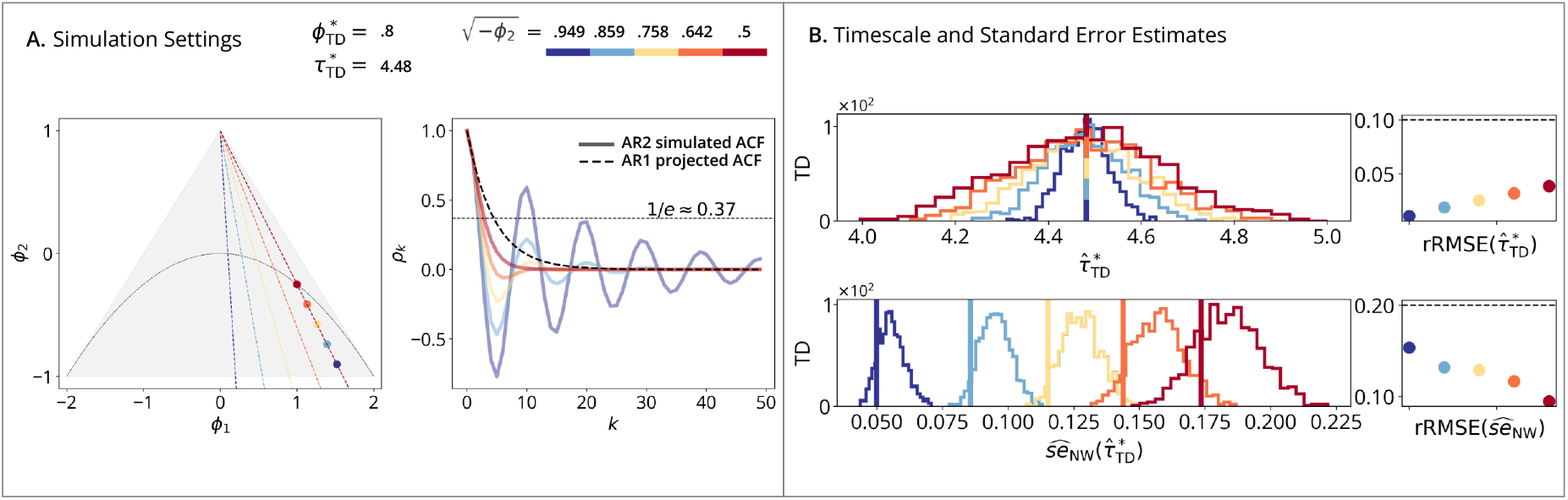
AR2 with damped oscillation simulations. (**A**) Simulation Setting (**Left**): triangle shows the AR2 stationary region in the (*ϕ*_1_, *ϕ*_2_) plane, with the periodic/aperiodic boundary at 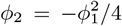 . Points show five selected AR2 (*ϕ*_1_, *ϕ*_2_) combinations that project to the same AR1 timescale, with increasing periodicity magnitude by adjusting 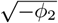 . (**Right**): solid lines show the simulated ACFs of the AR2 process defined in equation (19); the dashed black line shows the same AR1-projected ACF for all processes. (**B**) Timescale and Standard Error Estimates: vertical lines show the true timescale (*τ*_TD_ = 4.48 for all settings); histograms show estimates across *B* = 10, 000 replications; points show rRMSE. **(Row 1)** Time-Domain Linear Estimator, where rRMSE is plotted versus a 10% error line. (**Row 2**) Time-Domain Standard Error Estimator (Newey-West), where rRMSE is plotted versus a 20% error line.

Taken together, these results suggest that timescale inference can be reliably applied to oscillatory processes without explicitly disambiguating aperiodic and periodic components, a separation often required by frequency-domain methods (Donoghue et al., 2020; Gao et al., 2020). However, distinguishing neural from non-neural confounding remains critical for interpretation. We therefore recommend denoising methods such as RAPIDtide to remove confounding time-lagged oscillations, ensuring the estimated timescale more closely reflects neural rather than physiological processes (Frederick, 2025). That said, the maps reported in Data Analysis were robust to the exclusion of RAPIDtide denoising, suggesting the hierarchical organization of fMRI timescales is generalizable beyond this specific denoising strategy.

## 6 Conclusions

This study introduces statistical methods for mapping fMRI timescales, characterizing differences in local temporal autocorrelation across brain regions. We detail the large-sample properties of time- and autocorrelation-domain methods, showing that while both converge, they converge to different values due to their distinct parameter definitions. We also demonstrate that both estimators yield consistent standard errors under broad conditions, enabling reliable inference and hypothesis testing. This addresses a major limitation in fMRI timescale studies, which have typically reported point estimates without uncertainty measures.

Simulation results highlight differences in finite sample bias and variance between the methods. While all estimators are largely unbiased, the time-domain method performs as well as, or better than, the autocorrelation-domain method in terms of relative root mean square error (rRMSE). Standard error estimates exhibit pronounced differences between naive and Newey-West corrected methods, particularly under misspecified settings such as AR2 processes and realistic rfMRI data. Overall, the time-domain method (2) offers greater computational efficiency and estimation stability, whereas the autocorrelation-domain method (5) accommodates longer-range autocorrelations at the cost of reduced accuracy. The application of Newey-West corrected standard errors enhances inference reliability by reducing uncertainty bias across both methods (Newey and West, 1987), which is particularly vital when the autocorrelation decay is not strictly exponential.

Applied to HCP rfMRI data, both methods yield timescale and t-statistic maps consistent with known functional hierarchies (Van Essen et al., 2013). These results align with prior work demonstrating larger timescales in associative versus sensory cortices (Raut et al., 2020; Shafiei et al., 2020; Lurie et al., 2024; Mitra et al., 2014; Kaneoke et al., 2012; Wengler et al., 2020; Shinn et al., 2023; Manea et al., 2022; Ito et al., 2020; Müller et al., 2020). The spatial patterns reinforce the hypothesis that hierarchical organization governs temporal processing across cortical regions. However, mechanistic interpretations must carefully distinguish intrinsic neural dynamics from the confounding effects of hemodynamics and measurement noise. Linking macroscopic fMRI timescales to their underlying physiology remains an important objective for future research.

In conclusion, we present robust rfMRI estimators for timescales and standard errors, enabling rigorous statistical comparisons across regions, conditions, and subjects. These methods advance the accuracy and interpretability of neural timescale maps, moving beyond point estimates to incorporate uncertainty for inference and testing. This work lays the methodological foundation for future research into the role of timescales in brain structure and function.

## Appendix Autocorrelation-Domain Score Process

This appendix derives the score process *u*_*t*_ from equation (16). We omit scaling factors (e.g., 1/*K*), as they cancel out in the sandwich variance ratio 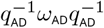 .

The pseudo-true parameter 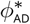 minimizes the population objective function *S*_*K*_(*ϕ*_AD_) (5), satisfying the first-order condition:

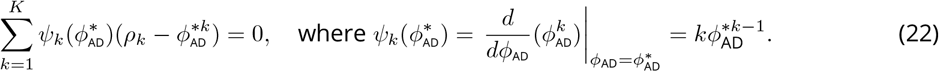

Using *ρ*_*k*_ = 𝔼[*X*_*t*_*X*_*t*−*k*_]/*γ*_0_ (1), we rewrite the residual term as an expectation:

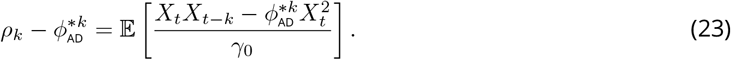

Substituting (23) into (22) defines the score process *u*_*t*_:

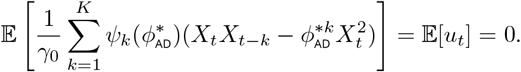

This matches the definition in equation (16). The asymptotic variance is then determined by the long-run variance of this score, 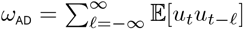 .

The sample score process *û*_*t*_ used for variance estimation is obtained by replacing population expectations with sample moments and the parameter 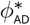 with the estimator 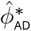 :

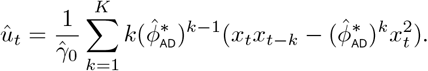

## Code and Data Availability

All simulation results and fMRI timescale maps, inclusive of the code by which they were derived, can be accessed on github.com/griegner/fmri-timescales. The code is under the open source MIT license, allowing access and reuse with attribution. The Human Connectome Project young adult dataset (ages 22-35; 2018 release) used in this study is publicly accessible under a data usage agreement, which describes specific terms for data use and sharing.

## Author Contributions

**Gabriel Riegner**: Conceptualization, Methodology, Software, Validation, Formal Analysis, Writing - Original Draft **Samuel Davenport**: Conceptualization, Methodology, Formal Analysis, Writing - Review & Editing, Supervision **Bradley Voytek**: Conceptualization, Writing - Review & Editing, Supervision **Armin Schwartzman**: Conceptualization, Methodology, Formal Analysis, Writing - Review & Editing, Supervision, Project Administration, Funding Acquisition.

